# A highly predictive signature of cognition and brain atrophy for progression to Alzheimer’s dementia

**DOI:** 10.1101/352344

**Authors:** Angela Tam, Christian Dansereau, Yasser Itturia-Medina, Sebastian Urchs, Pierre Orban, Hanad Sharmarke, John Breitner, Pierre Bellec, for the Alzheimer’s Disease Neuroimaging Initiative

## Abstract

Clinical trials in Alzheimer’s disease need to enroll patients whose cognition will decline over time, if left untreated, in order to demonstrate the efficacy of an intervention. Machine learning models used to screen for patients at risk of progression to dementia should therefore favor specificity (detecting only progressors) over sensitivity (detecting all progressors), especially when the prevalence of progressors is low. Here, we explore whether such high-risk patients can be identified using cognitive assessments and structural neuroimaging, by training machine learning tools in a high specificity regime. A multimodal signature of Alzheimer’s dementia was first extracted from ADNI1. We then validated the predictive value of this signature on ADNI1 patients with mild cognitive impairment (N=235). The signature was optimized to predict progression to dementia over three years with low sensitivity (55.1%) but high specificity (95.6%), resulting in only moderate accuracy (69.3%) but high positive predictive value (80.4%, adjusted for a “typical” 33% prevalence rate of true progressors). These results were replicated in ADNI2 (N=235), with 87.8% adjusted positive predictive value (96.7% specificity, 47.3% sensitivity, 85.1% accuracy). We found that cognitive measures alone could identify high-risk individuals, with structural measurements providing a slight improvement. The signature had comparable receiver operating characteristics to standard machine learning tools, yet a marked improvement in positive predictive value was achieved over the literature by selecting a high specificity operating point. The multimodal signature can be readily applied for the enrichment of clinical trials.

## Introduction

Alzheimer’s disease (AD), a leading cause of dementia, is marked by the abnormal accumulation of amyloid *β* (A*β*) and hyperphosphorylated tau proteins in the brain, which leads to widespread neurodegeneration. AD has a long prodromal phase, and it has been difficult to predict which individuals will decline and experience AD dementia. While mild cognitive impairment (MCI) puts individuals at risk, only a fraction (33.6% on average) of MCI patients will develop dementia within a period of three years, as shown in a meta-analysis of 41 studies [1]. Identifying MCI patients who will progress to AD dementia with enough specificity has thus been a challenge for clinical trials [2]. This lack of prognostic power may be due to individual variability. Different clinical phenotypes have been described where patients will exhibit distinct cognitive deficits [3]. Previous work has also characterized neuropathological subtypes based on the distribution of neurofibrillary tangles [4], which correspond well to distinct patterns of brain atrophy [5]. Different subtypes of brain atrophy have also been associated with different rates of progression to dementia [6]. The implications for prognosis are profound: only a subgroup of patients will experience a sharp cognitive decline that can be reliably predicted. We therefore propose to identify a subset of individuals with a homogenous signature of brain atrophy and cognitive deficits who will progress to AD dementia with high precision.

There is a large field focused on using machine learning to automatically detect MCI patients who will progress to AD dementia based on imaging and cognitive features. For models combining structural MRI and cognition, state-of-the-art performance is 79% accuracy (76% specificity, 83% sensitivity) [7]. Some groups have achieved higher accuracies ranging from 82-97% when using other imaging methods, such as A*β* positron emission tomography [8] or resting-state functional MRI [9]. Although this increase in accuracy may suggest that A*β* imaging and resting-state functional MRI are better features, these imaging measures are more invasive, costly, and currently lack the large scale of validation of tools that are already widely used in clinical settings, such as cognitive assessments and structural MRI. Given the need to develop tools that will easily scale up in clinical settings, we propose to focus on predictive models that use structural imaging and cognition as features.

Models are typically trained to maximize accuracy, defined as the proportion of subjects that were correctly identified, either as progressors or non-progressors. For enrichment in clinical trials, a more relevant metric is positive predictive value (PPV), which is the proportion of subjects that actually progress to dementia when they have been identified as such by the model. The PPV of a model is dependent on the baseline rate of progression in the sample, with a typical rate (within three years or more) in MCI patients being 33.6% [1]. Assuming a 33.6% baseline rate, it is possible to calculate the PPVs of published models in the literature, based on reported sensitivity and specificity scores. The adjusted PPV for models using cognitive and structural measures ranged from 50 to 75% [7,8,10–16]. In other words, up to half of subjects who were identified as progressors by published algorithms would not actually progress to dementia in a typical MCI sample. We therefore aimed to adapt the training regimen of predictive models to favor specificity over sensitivity, with the hypothesis that in this regime the models will identify progressors with high PPV. We expected that optimizing for high specificity will result in a low number of false positives, which is particularly important when the prevalence of progressors is low and therefore the susceptibility of the predictive model to identify false positive progressors is high.

The overall goal of this work was to develop a model to identify individuals who are at high risk of progression to AD dementia with high PPV and specificity, using structural MRI and cognitive features. We aimed to show that by training standard machine learning tools in a high specificity regime, we can identify the most robust progressor MCI patients with high confidence. We further wanted to assess whether those high risk individuals had prodromal AD, by examining longitudinal cognitive decline, as well as A*β* and tau burden in these individuals. We finally aimed to evaluate the complementarity of features derived from cognition and atrophy patterns by examining the overlap of high risk individuals who were identified as such by each modality. Although the complementarity of cognitive and structural measures has been extensively studied for prognosis of dementia in a general MCI population, the main contribution of this work is to examine their complementarity in the specific context of a high risk signature which achieves high specificity and PPV, at the cost of low sensitivity when the class of interest is relatively rare. Specific aims, as well as a summary of experiments and the main results, are listed in Table 1.

**Table 1.** Summary of objectives, experiments, and main findings

| Specific objectives | Experiments | Main findings |
| --- | --- | --- |
| 1) Identify subtypes of brain atrophy patterns | We used unsupervised clustering on atrophy maps generated from structural images in AD and CN subjects. | Seven distinct patterns of atrophy were identified, some of which were strongly associated with a diagnosis of AD (Figure 1b). |
| 2a) Replicate previous findings from works that used cognitive and structural features to predict progression to AD from MCI | A linear support vector machine, that was optimized for accuracy, was trained on the following features: 1) structural atrophy patterns, 2) multi-domain cognitive assessments, and 3) a combination of both. | The support vector machine based on cognitive features had higher predictive value than the structural MRI signature, similar to previous findings [7]. See Figures 2 and 3. |
| 2b) Train a model in a high specificity regime to identify high confidence AD subjects with a high-risk signature | We used a two-stage algorithm to ensure we were maximizing specificity over sensitivity. We trained on the following features: 1) structural atrophy patterns, 2) multi-domain cognitive assessments, and 3) a combination of both. | The two-stage algorithm resulted in a model that achieved high specificity and high PPV, with reduced sensitivity (Figure 2). Three high-risk signatures were generated (Figure 5). |
| 3) Assess if the high-risk signature generated by the two-stage algorithm can identify progressors in MCI subjects within a three year period | We measured PPV, specificity, sensitivity, and accuracy of the model in predicting progressors in two separate MCI cohorts. | The model achieved high specificity and high PPV, again at the cost of sensitivity and accuracy (Figures 2 and 4). |
| 4) Test the performance of the two-stage algorithm against standard algorithms | We compared the ROC performance of the two-stage algorithm against standard algorithms (e.g. KNN, GNB, SVM with a RBF kernel). | The performance of the two-stage algorithm did not differ from standard algorithms, in terms of area under a ROC curve, but was the only one to operate in a high-specificity, low sensitivity regime (Figure 3). |
| 5) Validate whether this high-risk signature represents a prodromal phase of AD | We compared cognitive decline, amyloid and tau burden in tagged high-risk individuals against those who were not. | Tagged high-risk individuals experienced sharper cognitive decline and higher levels of amyloid and tau than non-tagged individuals (Figure 4). |
| 6) Assess the complementarity of cognitive and structural measures | We examined whether there was overlap in the subjects that were identified by the three high-risk signatures. | The majority of subjects that were identified by the multimodal high-risk signature had been identified as such by the unimodal cognitive and unimodal structural signatures. The unimodal cognitive signature identified the majority of all high-risk subjects (Figure 6). |

## Materials and methods

### Data

Data used in the preparation of this article were obtained from the Alzheimer’s Disease Neuroimaging Initiative (ADNI) database (adni.loni.usc.edu). The ADNI was launched in 2003 as a public-private partnership, led by Principal Investigator Michael W. Weiner, MD. The primary goal of ADNI has been to test whether serial magnetic resonance imaging (MRI), positron emission tomography (PET), other biological markers, and clinical and neuropsychological assessment can be combined to measure the progression of mild cognitive impairment (MCI) and early Alzheimer’s disease (AD). For up-to-date information, see www.adni-info.org.

We took baseline T1-weighted MRI scans from the ADNI1 (228 CN, 397 MCI, 192 AD) and ADNI2 (218 CN, 354 MCI, 103 AD) studies. For a detailed description of MRI acquisition details, see http://adni.loni.usc.edu/methods/documents/mri-protocols/. All subjects gave informed consent to participate in these studies, which were approved by the research ethics committees of the institutions involved in data acquisition. Consent was obtained for data sharing and secondary analysis, the latter being approved by the ethics committee at the CRIUGM. For the MCI groups, each individual must have had at least 36 months of follow-up for inclusion in our analysis. We also further stratified the MCI groups into stable (sMCI), who never received any change in their diagnosis, and progressors (pMCI), who received a diagnosis of AD dementia within 36 months of follow-up. pMCI who progressed to AD dementia after 36 months were excluded. After applying these inclusion/exclusion criteria, we were left with 280 and 268 eligible MCI subjects in ADNI1 and ADNI2 respectively.

### Structural features from voxel-based morphometry

Images were processed with the NeuroImaging Analysis Kit (NIAK) version 0.18.1 (https://hub.docker.com/r/simexp/niak-boss/), the MINC toolkit (http://bic-mni.github.io/) version 0.3.18, and SPM12 (http://www.fil.ion.ucl.ac.uk/spm/software/spm12/) under CentOS with Octave (http://gnu.octave.org) version 4.0.2. Preprocessing of MRI data was executed in parallel on the Cedar supercomputer (https://docs.computecanada.ca/wiki/Cedar), using the Pipeline System for Octave and Matlab (PSOM) [21]. Each T1 image was linearly co-registered to the Montreal Neurological Institute (MNI) ICBM152 stereotaxic symmetric template [22], using the CIVET pipeline [23], and then re-oriented to the AC-PC line. Each image was segmented into grey matter, white matter, and CSF probabilistic maps. The DARTEL toolbox [24] was used to normalize the grey matter segmentations to a predefined grey matter template in MNI152 space. Each map was modulated to preserve the total amount of signal and smoothed with a 8 mm isotropic Gaussian blurring kernel. After quality control of the normalized grey matter segmentations, we were left with 621 subjects in ADNI1 (out of 700, 88.7% success rate) and 515 subjects in ADNI2 (out of 589, 87.4% success rate).

We extracted subtypes to characterize variability of grey matter distribution with the CN and AD samples from ADNI1. In order to reduce the impact of factors of no interest that may have influenced the clustering procedure, we regressed out age, sex, mean grey matter volume (GMV), and total intracranial volume (TIV), using a mass univariate linear regression model at each voxel. We then derived a spatial Pearson’s correlation coefficient between all pairs of individual maps after confound regression. This defined a subject x subject (377 × 377) similarity matrix which was entered into a Ward hierarchical clustering procedure (Figure 1a). Based on visual inspection of the similarity matrix, we identified 7 subgroups (Figure 1b). Each subtype was defined as the average map of each subgroup. For each subject, we computed spatial correlations between their map and each subtype, which we call weights (Figure 1a). The weights formed a n subject x m subtypes (n=377, m=7) matrix, which was included in the feature space for all predictive models including voxel-based morphometry (VBM) throughout this work. As in our previous works [20,25], we chose to use weights, which can be interpreted as continuous measures for subtype affinity, over discrete subtype membership because the latter is less informative as most individuals express similarity to multiple subtypes [26]. Note that although we chose to present our findings with 7 subtypes, we examined how the number of subtypes may impact our subsequent predictions. There was no significant difference in model performance when we changed the number of subtypes (see Table S1 in supplementary material).

**Figure 1.**
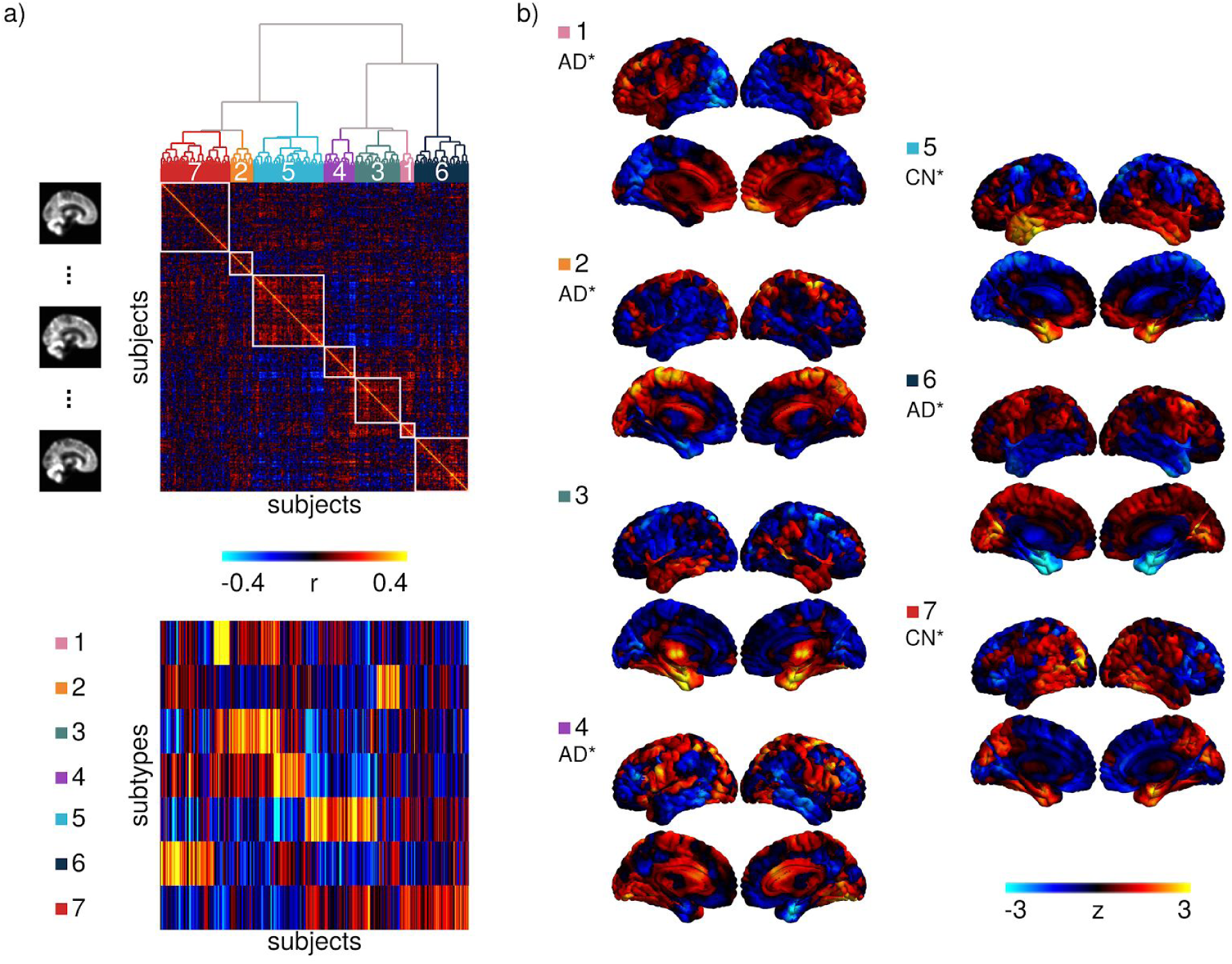
Subtyping procedure and resulting subtypes. a) A hierarchical clustering procedure identified 7 subtypes, or subgroups, of individuals with similar patterns of grey matter topography within the ADNI1 cohort of CN and AD subjects (top). A measure of spatial similarity, called subtype weight, between a single individual’s grey matter volume map and the average of a given subtype was calculated for all individuals and all subtypes (bottom). b) Maps of the 7 subtypes showing the distribution of grey matter across all voxels relative to the average. CN* and AD* denote significant associations between the subtype weights and diagnoses of cognitively normal (CN) or Alzheimer’s dementia (AD) respectively.

### Cognitive features

We took baseline neuropsychological scores for each subject from several cognitive domains: memory from the composite score ADNI-MEM [27], executive function from the composite score ADNI-EF [28], language from the Boston Naming Test (BNT), visuospatial from the clock drawing test, and global cognition from the Alzheimer’s Disease Assessment Scale-Cognitive (ADAS13). We chose measures that span multiple cognitive domains as it has been suggested that the use of a combination of neuropsychological measures is likely the best approach to predicting incipient dementia [29]. These scores were included as features for the predictive models involving cognition. Thirteen subjects across both ADNI1 and ADNI2 (8 AD, 5 MCI) had to be excluded due to missing values in their cognitive assessments. See Table 2 for demographic information of subjects who were included in analyses.

**Table 2.** Demographic information for post-QC subjects in ADNI1 and ADNI2.

| **ADNI1** | **CN** | **sMCI** | **pMCI** | **AD** |
| --- | --- | --- | --- | --- |
| **N** | 205 | 88 | 147 | 165 |
| **Age ± SD** | 76.1 ± 5.0 | 74.0 ± 7.6 | 74.3 ± 7.1 | 75.4 ± 7.5 |
| **Female %** | 51.7 | 40.9 | 40.8 | 51.5 |
| **APOE4+ %** | 27.8 | 37.5 | 68.7 | 65.4 |
| **ADAS13 ± SD** | 9.5 ± 4.3 | 14.3 ± 5.5 | 21.3 ± 5.3 | 28.6 ± 7.1 |
| **MMSE ± SD** | 29.1 ± 1.0 | 27.7 ± 1.7 | 26.7 ± 1.7 | 23.4 ± 2.0 |
| **ADNI2** | **CN** | **sMCI** | **pMCI** | **AD** |
| **N** | 188 | 180 | 55 | 89 |
| **Age ± SD** | 72.8 ± 6.1 | 70.8 ± 7.3 | 72.1 ± 7.1 | 74.4 ± 7.8 |
| **Female %** | 54.0 | 47.8 | 49.1 | 46.1 |
| **APOE4+ %** | 29.4 | 35.6 | 65.4 | 71.3 |
| **ADAS13 ± SD** | 9.1 ± 4.2 | 11.8 ± 5.3 | 21.4 ± 6.5 | 31.6 ± 8.7 |
| **MMSE ± SD** | 29.1 ± 1.1 | 28.4 ± 1.6 | 27.3 ± 1.9 | 23.1 ± 2.3 |

ADAS13=Alzheimer's Disease Assessment Scale - Cognitive subscale (13 items);
MMSE=Mini Mental State Examination

### Prediction of high confidence AD dementia cases in ADNI1

We trained a linear support vector machine (SVM) model with a linear kernel, as implemented by Scikit-learn [30] version 0.18 to classify AD vs CN from ADNI1 to get a baseline prediction accuracy. We then used a two-step method to select an operating point for the linear SVM to obtain a highly precise and specific classification [20]. This was done by replicating the SVM prediction via subsampling and identifying the patients with highly robust prediction outcomes, i.e. who are consistently identified as true AD cases (true positives) during testing, regardless of the training subsample. This approach was found, in practice, to lead to a highly specific prediction, in addition to offering a guarantee of robustness; see [20] for more information. Specifically here, a tenfold cross-validation loop was used to estimate the performance of the trained model. Classes were balanced inversely proportional to class frequencies in the input data for the training. A nested cross-validation loop (stratified shuffle split with 50 splits and 20% test size, i.e. a random permutation cross-validator was used to split the data into 50 training and test sets, where the size of the test set was always 20% of the original sample size) was used for the grid search of the SVM hyperparameter *C* (grid was 10^−2^ to 10^1^ with 15 equal steps). We randomly selected subsamples of the dataset, retaining a set percentage of participants in each subsample. For each subsample, a separate SVM model was trained to predict AD or CN in ADNI1. The SVM training was replicated a number of times. Both the subsample size and the number of subsamples were selected to maximize the positive predictive value of the prediction of sMCI vs pMCI in ADNI1, as described below. Predictions were made on the remaining subjects that were not used for training, and, for each subject, we calculated a hit probability defined as the frequency of correct classification across all SVM replications in which the test set contained that subject. High confidence AD cases were defined as individuals with 100% hit probabilities with the AD label. Next, we trained a logistic regression classifier [31], with L1 regularization on the coefficients, to predict the high confidence AD cases. A stratified shuffle split (500 splits, 50% test size) was used to estimate the performance of the model for the grid search of the hyperparameter *C* (grid was 10^−2^ to 10^1^ with 15 equal steps) on the overall ADNI1 sample, and the same hyperparameters were used for all SVM replications.

We used the entire CN and AD sample from ADNI1 to obtain three highly predictive signatures (HPS) (i.e. models), 1) one using VBM subtype weights as features (VBM only), 2) one using only cognitive features (COG only), 3) and one using the combination of VBM subtype weights and cognitive features (VCOG). In all three signatures, age, sex, mean GMV, and TIV were also included as features.

### Prediction of progression to AD dementia from the MCI stage in ADNI1

The logistic regression trained on AD vs CN was used to identify MCI patients who have a HPS of AD dementia in ADNI1. Our hyperparameters for this logistic regression were optimized based on the number of subsamples and subsample size that produced the maximum specificity and PPV for the classification of sMCI (n=89) vs pMCI (n=155) in ADNI1, while maintaining a minimum of 30% sensitivity. We varied the number of subsamples (100, 500, 1000) and subsample size (10%, 20%, 30%, 50%) to perturb the model and identify subjects that had robust outcomes during the testing phase regardless of the training subsample. We then re-trained our models to classify AD vs CN in ADNI1 with these optimized hyperparameters. This was done for all three signatures. In brief, we used the AD and CN sample from ADNI1 as a training set, and the MCI subjects from ADNI1 as a validation set. The ADNI2 sample was then used as an independent replication (test) set, to establish the performance of the two-stage model without further changes to the hyperparameters.

### Statistical test of differences in model performance

We used Monte-Carlo simulations to generate confidence intervals on the performance (i.e. accuracy, PPV, specificity and sensitivity) of both linear SVM and HPS models for their predictions of AD vs CN and pMCI vs sMCI. Taking the observed sensitivity and specificity, and using similar sample sizes to our experiment, we replicated the number of true and false positive detection 100000 times using independent Bernoulli variables, and derived replications of PPV, specificity and sensitivity. By comparing these replications to the accuracy, sensitivity, specificity and PPV observed in both models, we estimated a p-value for differences in model performance [32]. A p-value smaller than 0.05 was interpreted as evidence of a significant difference in performance, and 0.001 as strong evidence. We also used this approach to compare the performance of the combined features (VCOG) to the models containing VBM features (VBM) or cognitive features (COG) only. Note that, based on our hypotheses regarding the behaviour of the HPS model, the tests were one-sided for increased accuracy, specificity and PPV, and one-sided for decreased sensitivity.

To assess the performance of the HPS models against standard machine learning algorithms, we used four algorithms (SVM with a RBF kernel, K nearest neighbors, random forest, and Gaussian naive Bayes) to train models to classify AD vs CN in the ADNI1 dataset. We then tested and validated these models on classifying AD vs CN in ADNI2 and finally pMCI vs sMCI in both ADNI1 and ADNI2 separately. See the supplementary material for details of the implementations of these latter algorithms. We then generated ROC curves and calculated the area under the curve (AUC) for each model and classification (AD vs CN; pMCI vs sMCI) in both ADNI1 and ADNI2.

### Statistical tests of association of progression, AD biomarkers, and risk factors in high confidence MCI subjects

Based on the classifications resulting from the linear SVM and HPS models, we separated the MCI subjects into three different groups: 1) High confidence, subjects who were selected by the HPS model as hits, 2) Low confidence, subjects who were selected by the linear SVM model as hits but were not selected by the HPS model, and 3) Negative, subjects who were not selected as hits by either algorithm.

In order to validate whether the high confidence subjects represented individuals who were in a prodromal phase of AD, we tested if this subgroup was enriched for progression to dementia, APOE4 carriers, females, and subjects who were positive for A*β* and tau pathology. Positivity of AD pathology was determined by CSF measurements of A*β* 1-42 peptide and total tau with cut-off values of less than 192 pg/mL and greater than 93 pg/mL respectively [33]. We implemented Monte-Carlo simulations, where we selected 100000 random subgroups out of the original MCI sample. By comparing the proportion of progressors, APOE4 carriers, females, A*β*-positive, and tau-positive subjects in these null replications to the actual observed values in the HPS subgroup, we estimated a p-value [32] (one sided for increase). A p-value smaller than 0.05 was interpreted as evidence of a significant enrichment, and 0.001 as strong evidence.

One-way ANOVAs were used to evaluate differences between the HPS groupings with respect to age. Post-hoc Tukey’s HSD tests were done to assess pairwise differences among the three classes (high confidence, low confidence, negative). These tests were implemented in Python with the SciPy library [34] version 0.19.1 and StatsModels library [35] version 0.8.0.

To explore the impact of HPS grouping on cognitive trajectories, linear mixed effects models were performed to evaluate the main effects of and interactions between the HPS groups and time on ADAS13 scores up to 36 months of follow-up. The models were first fit with a random effect of participant and then were fit with random slopes (time | participant) if ANOVAs comparing the likelihood ratio suggested a significant improvement in model fit. All tests were performed separately on the ADNI1 and ADNI2 datasets. These tests were implemented in R version 3.3.2 with the library nlme version 3.1.128 [36].

### Public code, data availability and reproducibility

The code used in this experiment is available on a GitHub repository (https://github.com/SIMEXP/vcog_hps_ad) and zenodo (https://doi.org/10.5281/zenodo.1444081).

We shared a notebook replicating all the machine learning experiments, starting after the generation of VBM subtypes. However, in order to protect the privacy of the study participants, we could not share individual subtype weights alongside other behavioural data and diagnostic information. We thus created parametric (Gaussian) bootstrap simulations, based on group statistics alone, that will allow interested readers to replicate results similar to those presented in this manuscript, using the exact same code and computational environment that were used on real data, but with purely synthetic data instead. The notebook can be executed online via the binder platform (http://mybinder.org), and runs into a docker container (https://mybinder.org/v2/gh/SIMEXP/vcog_hps_ad/master?filepath=%2Fvcog_hpc_prediction_simulated_data.ipynb), built from a configuration file that is available on GitHub (https://github.com/SIMEXP/vcog_hps_ad/blob/master/Dockerfile). The container itself is available on Docker Hub (https://hub.docker.com/r/simexp/vcog_hps_ad/). The simulated data was archived on figshare (https://figshare.com/articles/Simulated_cognitive_and_structural_MRI_data_from_ADNI/7132757).

The simulation included the following 16 variables: age, sex, mean grey matter volume, total intracranial volume, 5 cognitive assessment scores and 7 VBM subtype weights from both ADNI1 and ADNI2. Subjects that had missing values for these variables were discarded from the simulation, leaving N=1115 subjects. We stratified the population into 12 subgroups: the four clinical labels (AD, pMCI, sMCI, CN), each further subdivided by the three prediction subclasses identified in this paper (negative, low confidence, high confidence). For each subgroup, we estimated the average and covariance matrices between the 16 variables of interest. We then generated a number of multivariate normal data points that matched the number of subjects found in each subgroup, using the empirical mean and covariance matrix of each subgroup. Finally, the range of the simulated data was clipped to the range of the original real data, and the simulated sex data points were binarized by nearest neighbour.

The statistics from the predictive model in the original implementation are similar to that of the simulated data. The model predicted the progression of dementia from MCI in ADNI1 with a PPV of 93.1% (specificity of 93.2%) on real data. This coincides with a 93.3% PPV (specificity of 94.3%) that we get when using the simulated data. Similarly, with the ADNI2 dataset the model achieved a 81.3% PPV (specificity of 96.7%) from the real data and a 75.7% PPV (specificity of 95.0%) from the simulated data.

## Results

### Subtypes of brain atrophy

Subtype 1 was characterized by reduced relative GMV in the occipital, parietal and posterior temporal lobes. Subtype 2 displayed reduced relative GMV across the cortex, except for the medial parts of the parietal and occipital lobes and the cingulate. Subtype 3 had increased relative GMV in the medial and lateral temporal lobes, insula, and striatum. Subtype 4 had decreased relative GMV in the temporal lobes, inferior parietal lobes, posterior cingulate, and the prefrontal cortices. Subtype 5 was characterized by greater relative GMV in the temporal lobes, while Subtype 6 had the opposite pattern of reduced relative GMV in the temporal lobes. Subtype 7 displayed greater relative GMV in the parietal lobes, posterior lateral temporal lobes, medial temporal lobes, and medial occipital lobes. See Figure 1b for surface representations of the subtypes. Diagnosis (CN, sMCI, pMCI, AD) accounted for a substantial amount of variance in subtype weights for subtypes 1 (F=8.51, p=1.30 × 10^−5^), 2 (F=10.32, p=1.00 × 10^−6^), 4 (F=14.53, p=2.60 × 10^−9^), 5 (F=13.86, p=6.77 × 10^−9^), 6 (F=34.27, p=2.57 × 10^−21^), and 7 (F=37.02, p=5.85 × 10^−23^). Post-hoc t-tests showed AD subjects had significantly higher weights compared to CN (Figure 1b) for subtypes 1 (t=2.88, p=0.02), 2 (t=4.05, p=3.0 × 10^−4^), 4 (t=4.83, p<1.0 × 10^−4^), and 6 (t=7.86, p=<1.0 10^−4^), making these subtypes associated with a diagnosis of AD. CN subjects had significantly higher weights compared to AD for subtypes 5 (t=−4.86, p<1.0 × 10^−4^) and 7 (t=−6.95, p<1.0× 10^−4^), making these subtypes associated with a cognitively normal status.

### Prediction of AD dementia vs cognitively normal individuals

The linear SVM model trained using the VCOG features achieved 94.5% PPV (95.6% specificity, 93.9% sensitivity, 94.9% accuracy) when classifying AD vs CN in ADNI1. Such high performance was expected given the marked cognitive deficits associated with clinical dementia. COG features only actually reached excellent performance as well (97.6% PPV, 98.0% specificity, 96.4% sensitivity, 97.3% accuracy), while using VBM features only yielded markedly lower performances (86.4% PPV, 89.3% specificity, 79.6% sensitivity, 84.8% accuracy) (see Figures 2 and ROC analysis in Figure 3). Note that the performance metrics in ADNI1 were estimated through cross-validation, and represent an average performance for several models trained on different subsets of ADNI1. We then trained a model on all of ADNI1, and estimated its performance on an independent dataset, ADNI2. Using VCOG predictors, the ADNI1 model reached 92.0% PPV (96.3% specificity, 92.0% sensitivity, 94.5% accuracy), when applied on ADNI2 AD vs CN data. Again the performance was comparable with COG predictors only (92.2% PPV, 96.3% specificity, 94.3% sensitivity, 95.6% accuracy), and VBM features only achieved lower performance (57.3% PPV, 79.8% specificity, 56.7% sensitivity, 72.3% accuracy) (see Figures 2 and ROC analysis in Figure 3). Note that PPV is dependent on the proportion of patients and controls for a given sensitivity and specificity. Since the ADNI2 sample had a substantially smaller proportion of AD subjects compared to ADNI1, the resulting PPV was reduced. When we adjusted the baseline rate of AD subjects in ADNI2 to the same rate in ADNI1, the PPVs were 95.2%, 95.3%, and 70.2% for the VCOG, COG, and VBM models respectively.

**Figure 2.**
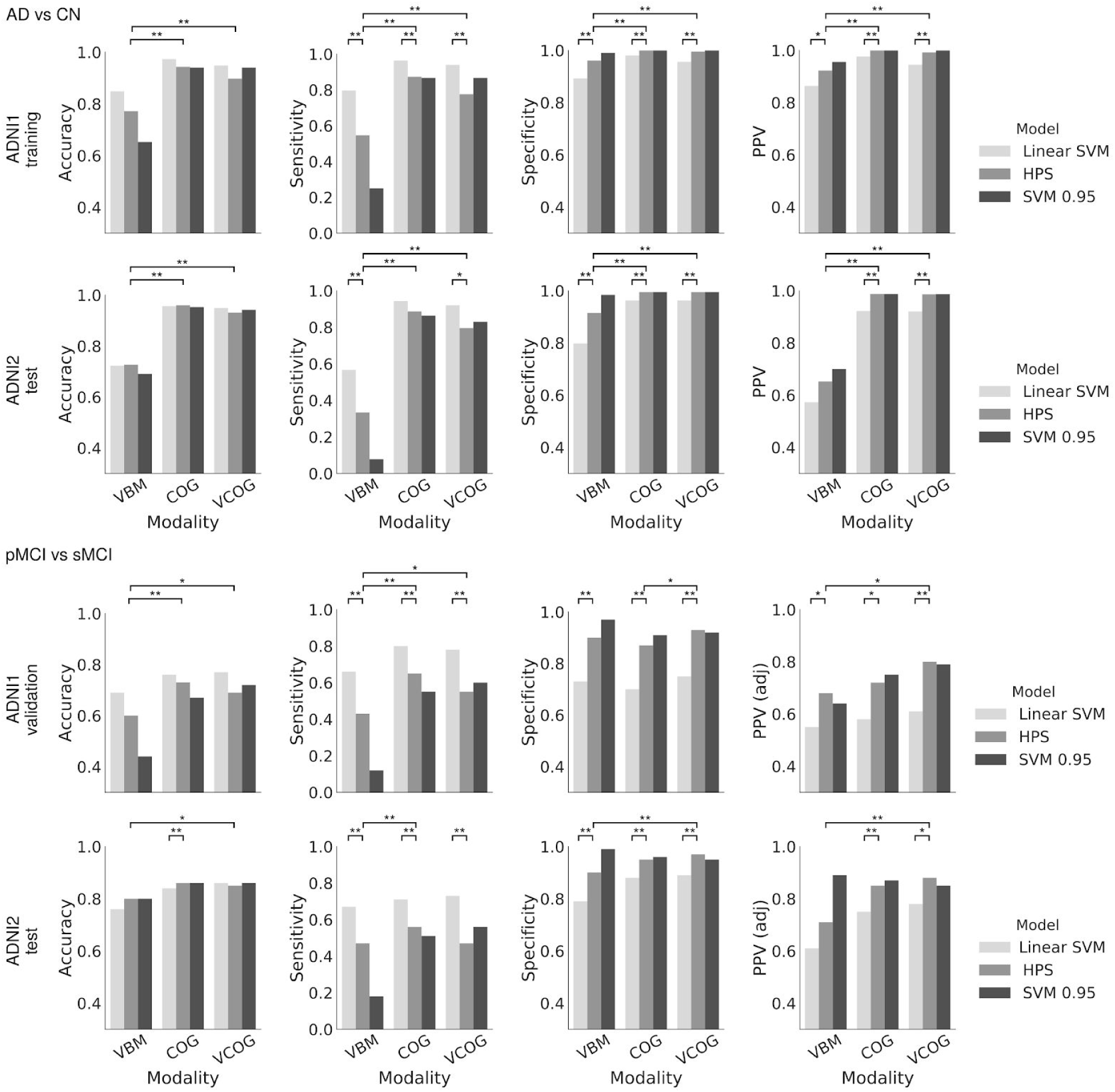
Accuracy, specificity, sensitivity, and positive predictive value (PPV) for different classifiers: linear SVM, highly predictive signature (HPS), and the linear SVM thresholded at 0.95 (SVM 0.95), for the classifications of patients with AD dementia (AD) and cognitively normal individuals (CN) and patients with mild cognitive impairment who progress to AD (pMCI) and stable MCI (sMCI) in ADNI1 and ADNI2. VBM represents the model trained with VBM subtypes, COG represents the model trained with baseline cognitive scores, and VCOG represents the model trained with both VBM subtypes and cognition. Positive predictive value was adjusted (PPV (adj)) for a prevalence of 33.6% pMCI in a sample of MCI subjects for both ADNI1 and ADNI2 MCI cohorts. Significant differences are denoted by * for p<0.05 and ** for p<0.001).

**Figure 3.**
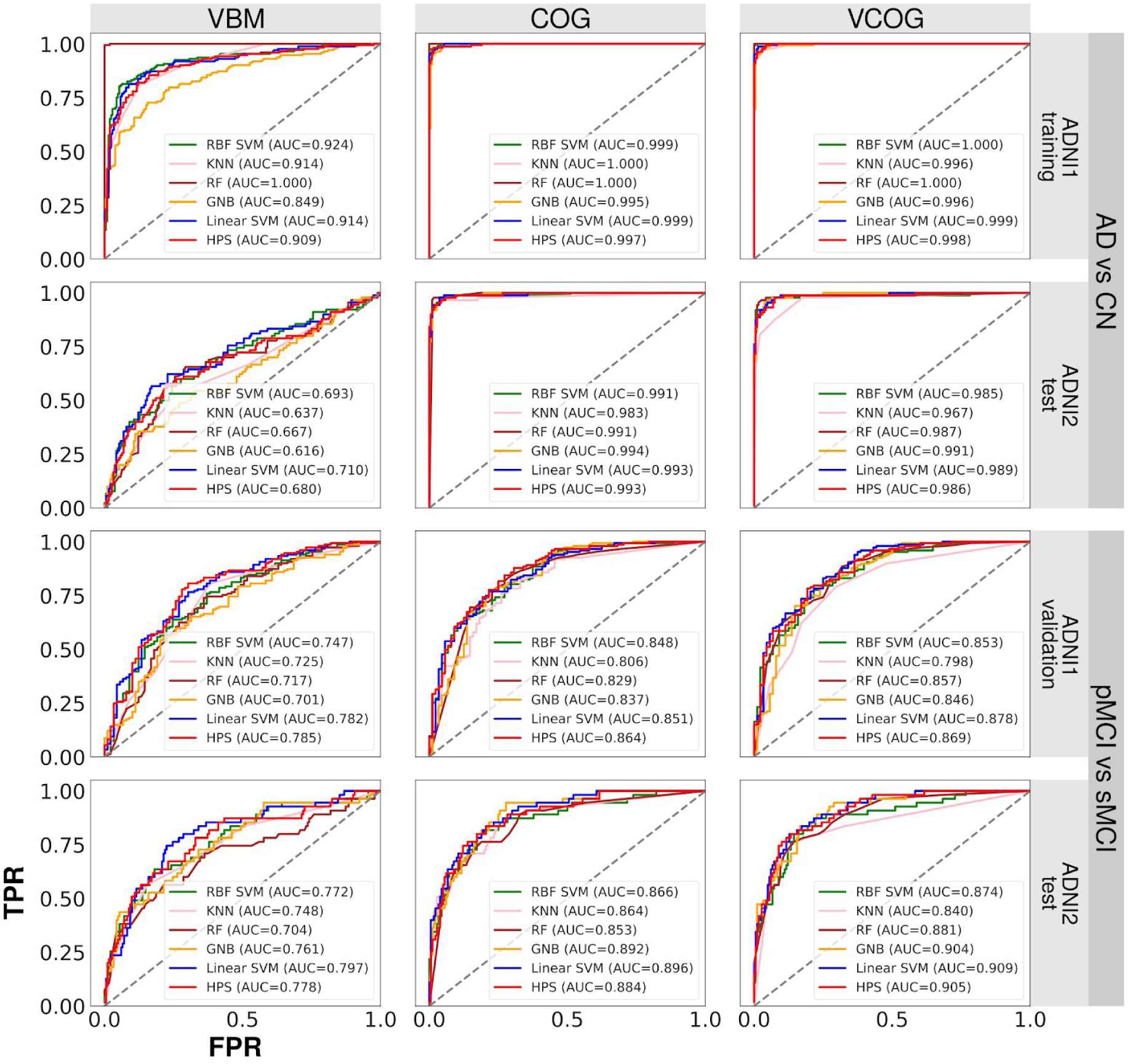
Receiver operating characteristic (ROC) curves for various machine learning algorithms with different features (VBM for VBM subtypes only, COG for cognitive features only, VCOG for a combination of VBM subtypes and cognitive features). Algorithms included a support vector machine with a radial basis function kernel (RBF SVM), K nearest neighbors (KNN), random forest (RF), Gaussian naive Bayes (GNB), a support vector machine with a linear kernel representing the first stage (Linear SVM) of the two-stage predictive model, and the two-stage highly predictive signature (HPS). TPR refers to true positive rate, FPR refers to false positive rate, and AUC refers to area under the curve.

### Identification of high confidence AD cases for prediction

The VCOG HPS model achieved 99.2% PPV (99.5% specificity, 77.6% sensitivity, 89.7% accuracy) in classifying high confidence AD subjects in ADNI1. These performance scores were estimated by cross-validation of the entire two-stage process (training of SVM, estimation of hit probability, identification of HPS). However, the hyperparameters of the two-stage model were optimized on classifying pMCI vs sMCI in ADNI1, as described previously. We next trained a single model on all of ADNI1, which we applied on an independent sample (ADNI2). The ADNI1 AD VCOG HPS model reached 98.6% PPV (99.5% specificity, 79.5% sensitivity, 93.1% accuracy) on ADNI2. As was previously observed with the conventional SVM analysis, the VCOG HPS model had similar performance to the COG HPS model (ADNI1: 100% PPV, 100% specificity, 87.3% sensitivity, 94.2% accuracy; ADNI2: 98.7% PPV, 99.5% specificity, 88.6% sensitivity, 96.0% accuracy), and outperformed the VBM HPS model (ADNI1: 92.3% PPV, 96.1% specificity, 54.6% sensitivity, 77.2% accuracy; ADNI2: 65.2% PPV, 91.5% specificity, 33.3% sensitivity, 72.7% accuracy); see Figure 2. When adjusted to the same baseline rate of AD subjects as ADNI1, the PPVs in ADNI2 were 99.2%, 99.3%, and 76.7% for the VCOG, COG, and VBM HPS models respectively.

### High confidence prediction of progression to AD dementia

When using the full VCOG features, 87 MCI patients were selected as high confidence in ADNI1, out of which 81 (93.1% PPV) were pMCI within 36 months of follow-up. This represented a large, significant increase over the baseline rate of progressors in the entire ADNI1 MCI sample (37.4%) (p<0.001). This was also a significant increase over the SVM’s predictions, where 83.9% of subjects that it had labeled as hits were true progressors (p<0.001). When adjusted to a 33.6% baseline rate of progressors, more typical of MCI populations, the PPV of high confidence subjects for prognosis of dementia was 80.4% (93.2% specificity, 55.1% sensitivity, 69.3% accuracy).

We replicated these analyses in the MCI sample from ADNI2 (N=235). Using VCOG features, 32 subjects were identified as high confidence, 26 of which progressed to AD dementia within 36 months follow-up (81.2% PPV, specificity of 96.7%, sensitivity of 47.3%, 85.1% accuracy, 87.8% PPV adjusted to a 33.6% baseline rate). This represented a significantly higher prevalence than the 30.6% baseline rate in the entire ADNI2 MCI cohort (p<0.001). This was also a significant increase over the SVM’s predictions, where 67.8% of subjects it had labeled as hits were true progressors (p<0.001).

As in the classifications of AD vs CN, the VCOG HPS model tended to have higher performance compared to the VBM HPS (ADNI1: 89.9% specificity, 42.9% sensitivity, 60.5% accuracy, 87.7% PPV, 68.2% adjusted PPV; ADNI2: 90.1% specificity, 47.3% sensitivity, 80.2% accuracy, 59.1% PPV, 70.7% adjusted PPV) in classifying pMCI vs sMCI. The VCOG HPS also had similar performance compared to the COG HPS (ADNI1: 87.5% specificity, 64.6% sensitivity, 73.2% accuracy, 89.6% PPV, 72.3% adjusted PPV; ADNI2: 95.0% specificity, 56.4% sensitivity, 86.0% accuracy, 77.5% PPV, 85.1% adjusted PPV) for distinguishing between pMCI and sMCI. Notably, the VCOG features lead to higher PPV than VBM and COG features taken independently, both in ADNI1 and ADNI2. That increase was large and significant between VCOG and VBM (up to 17%) and marginal and non-significant between VCOG and COG (up to 8%); see Figure 2.

### Trade-off between sensitivity and specificity of different algorithms

The HPS models consistently outperformed the linear SVM classifiers with respect to specificity (p<0.001) in the classifications of AD vs CN and pMCI vs sMCI in both ADNI1 and ADNI2, regardless of the features that the models contained. The HPS also had greater PPV (p<0.05) adjusted for a typical prevalence of 33.6% pMCI in a given sample of MCI subjects [1]. However, these increases in specificity and PPV for the HPS model came at a significant cost of reduced sensitivity compared to the linear SVM classifier, across all models in both ADNI1 and ADNI2 (p<0.05) (Figure 2). Note that this shift towards lower sensitivity and higher specificity and PPV could be achieved by adjusting the threshold of the SVM analysis (see Figure 2 and ROC analysis in Figure 3), and is not unique to the two-stage procedure we implemented. This trade-off between sensitivity and specificity is universal across machine learning algorithms and similar results can be achieved by adjusting the prediction threshold of different strategies. As shown by the ROC curves and AUC values in Figure 3, other machine learning algorithms (SVM with a radial basis function kernel, K nearest neighbors, random forest, and Gaussian naive Bayes) also performed similarly to the HPS. Thus, the value of the HPS is in the selection of a threshold point in order to operate in a high specificity regime.

### Characteristics of MCI subjects with a highly predictive VCOG signature of AD

High confidence MCI subjects with the VCOG signature were more likely to be progressors (Figure 4a) compared to low confidence subjects and negative subjects (ADNI1: p<0.001; ADNI2: p<0.001). High confidence MCI subjects were also more likely to be APOE4 carriers (Figure 4b) (ADNI1: p<0.005; ADNI2: p<0.05). There was no difference in sex across the HPS groupings in the MCI subjects of either the ADNI1 or ADNI2 cohorts (Figure 4c). This was consistent with the whole sample, where there were equal proportions of progressors across both sexes in each dataset (ADNI1: χ^2^=0.015, p=0.90; ADNI2: χ^2^=0.0002, p=0.99). The high confidence class was also significantly enriched for A*β*-positive subjects in ADNI1 (p<0.05). However, this result was not replicated in the ADNI2 MCI subjects (Figure 4d). Similarly with tau, we found a significant increase in tau-positive subjects in the high confidence group of ADNI1 (p<0.05), but not in ADNI2 (Figure 4e). We found a significant age difference across the HPS classes in ADNI2 (F=5.68, p<0.005), where the high confidence subjects were older than the Negative subjects by a mean of 4.4 years. However, age did not differ across the HPS classes in ADNI1 (Figure 4f). Finally, high confidence subjects had significantly steeper cognitive declines compared to the low confidence and negative groups (Figure 4g): there were significant interactions between the HPS groupings and time in ADNI1: (high confidence *β*=−0.147, t=−7.56, p<0.001; low confidence *β*=−0.055, t=−2.46, p<0.05) and ADNI2 (high confidence *β*=−0.194, t=−8.69, p<0.001; low confidence *β*=−0.072, t=−3.31, p=0.001). The high confidence subjects in ADNI1 and ADNI2 respectively gained 1.8 and 2.3 more points each year on the ADAS13 compared to the low confidence and negative groups. Note that higher scores on the Alzheimer’s Disease Assessment Scale - Cognitive subscale (13 items) (ADAS13) represent worse cognitive function.

**Figure 4.**
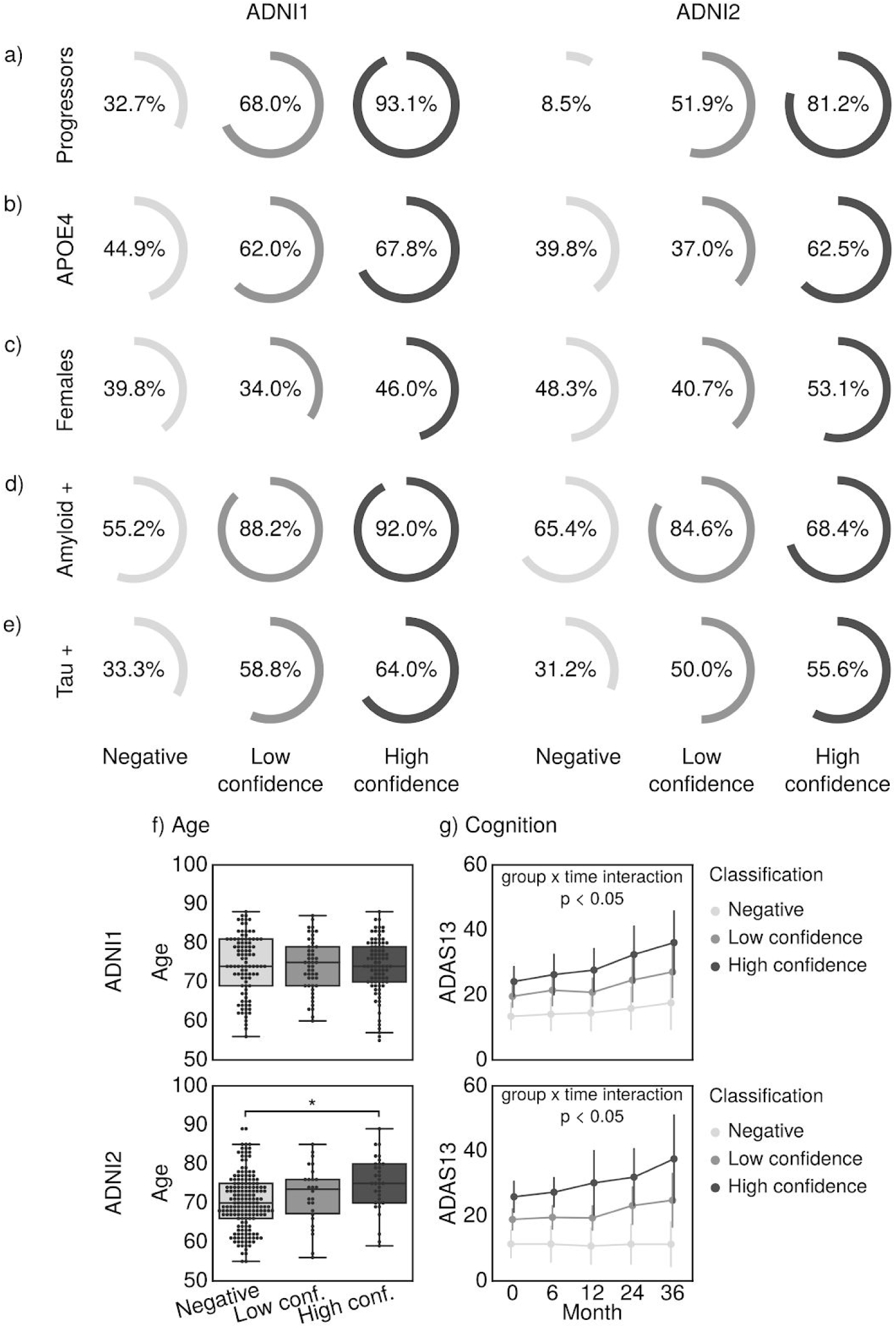
Characteristics of MCI subjects with the VCOG signature in ADNI1 and ADNI2. We show the percentage of MCI subjects who a) progressed to dementia, were b) APOE4 carriers, c) female, d) positive for A*β* measured by a cut-off of 192 pg/mL in the CSF [22], and e) positive for tau measured by a cut-off of 93 pg/mL in the CSF [22] in each classification (High confidence, Low confidence, and Negative). f) Age and g) cognitive trajectories, measured by the Alzheimer’s Disease Assessment Scale - Cognitive subscale with 13 items (ADAS13), across the three classes. Significant differences are denoted by * for family-wise error rate-corrected p<0.05.

### COG, VBM and VCOG highly predictive signatures

The COG signature was mainly driven by scores from the ADAS13, which measures overall cognition, ADNI-MEM, a composite score that measures memory [27], and ADNI-EF, a composite score that measures executive function [37] (coefficients were 5.49, −4.80 and −2.50 respectively). In this model, sex, age, mean GMV, and TIV contributed very little, relative to the cognitive features (Figure 5b). Note that these coefficients should be interpreted as pseudo z-scores as the features had been normalized to zero mean and unit variance.

**Figure 5.**
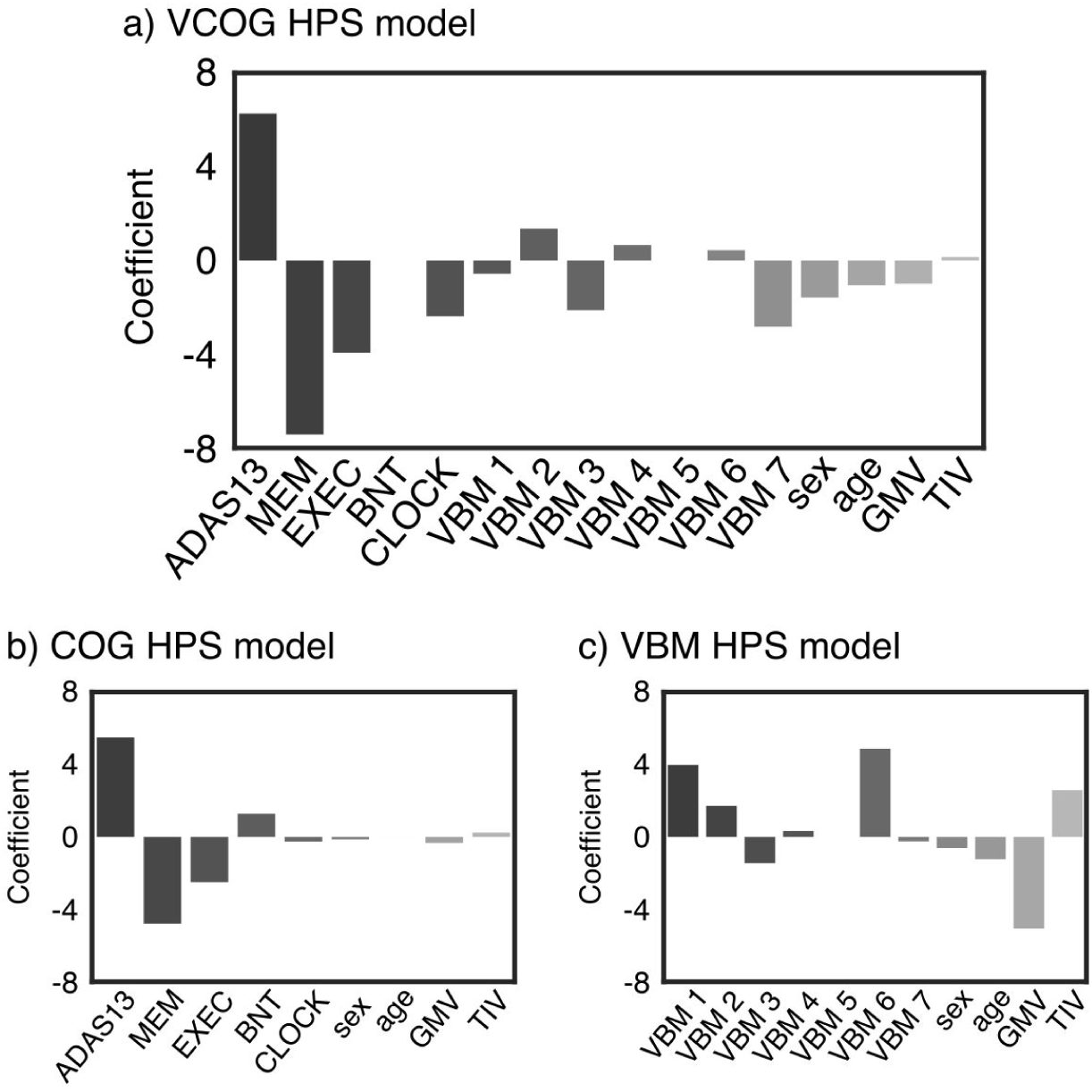
Coefficients of the high confidence prediction a) VCOG HPS model, b) COG HPS model, c) VBM HPS model. ADAS13=Alzheimer’s Disease Assessment Scale - Cognitive, MEM=ADNI-MEM score; EXEC=ADNI-EF score, BNT=Boston Naming Test, CLOCK=clock drawing test, VBM 1-7=VBM subtype weights, GMV=mean grey matter volume, TIV=total intracranial volume.

Almost all grey matter subtypes contributed to the VBM signature. Mean GMV, subtype 1 (reduced relative GMV in the occipital, parietal and posterior temporal lobes) and subtype 6 (reduced relative GMV in the temporal lobes, notably the medial temporal regions) had the highest weights in the model (coefficients were −5.07, 4.87, and 3.98 respectively) (Figure 5c). We had anticipated the larger contribution of these two subtypes as they have been described in previous AD subtyping work [5,17–19].

The ADAS13, memory (ADNI-MEM) and executive function (ADNI-EF) scores contributed the most to the VCOG signature (coefficients were 6.27, −7.43 and −3.95 respectively, Figure 5a). Of the VBM features, subtypes 2, 3 and 7 contributed the most to the signature (coefficients were 1.36, −2.12 and −2.83 respectively). Subtypes 1 and 6, which had the highest positive weights in the VBM HPS model, were given marginal weights in the VCOG HPS model, which is potentially indicative of redundancy with COG features. Note that the weights for subtypes 3 and 7 were negative in the model, which means that predicted AD and pMCI cases had brain atrophy patterns that were spatially dissimilar to those subtypes.

### Comparison of COG, VBM and VCOG high confidence subjects

We found substantial overlap of subjects labeled as high confidence in the MCI cohorts across the VBM, COG and VCOG signatures (Figure 6). There were very few subjects that were labeled as high confidence exclusively by the VCOG signature. As to be expected, the majority of subjects labeled as high confidence by the VCOG signature (ADNI1: 97.7%; ADNI2: 100%) were also labeled as high confidence by either the VBM only or COG only signatures or both. Of the subjects that were labeled as high confidence by the VBM only signature, 23.6% and 55.2% in ADNI1 and ADNI2 respectively were identified exclusively by the VBM HPS. There were relatively few subjects (7 and 2 subjects in ADNI1 and ADNI2 respectively) that were captured by VBM and VCOG but missed by the COG HPS. The COG HPS actually identified the majority of all high confidence subjects across the three signatures (ADNI1: 106 of 132 total subjects, ADNI2: 40 of 65 total subjects). From Figure 6, we can see that the VCOG HPS acts as a refinement of the COG signature, as the VCOG HPS captures a subset of subjects that were labeled by the COG HPS.

**Figure 6.**
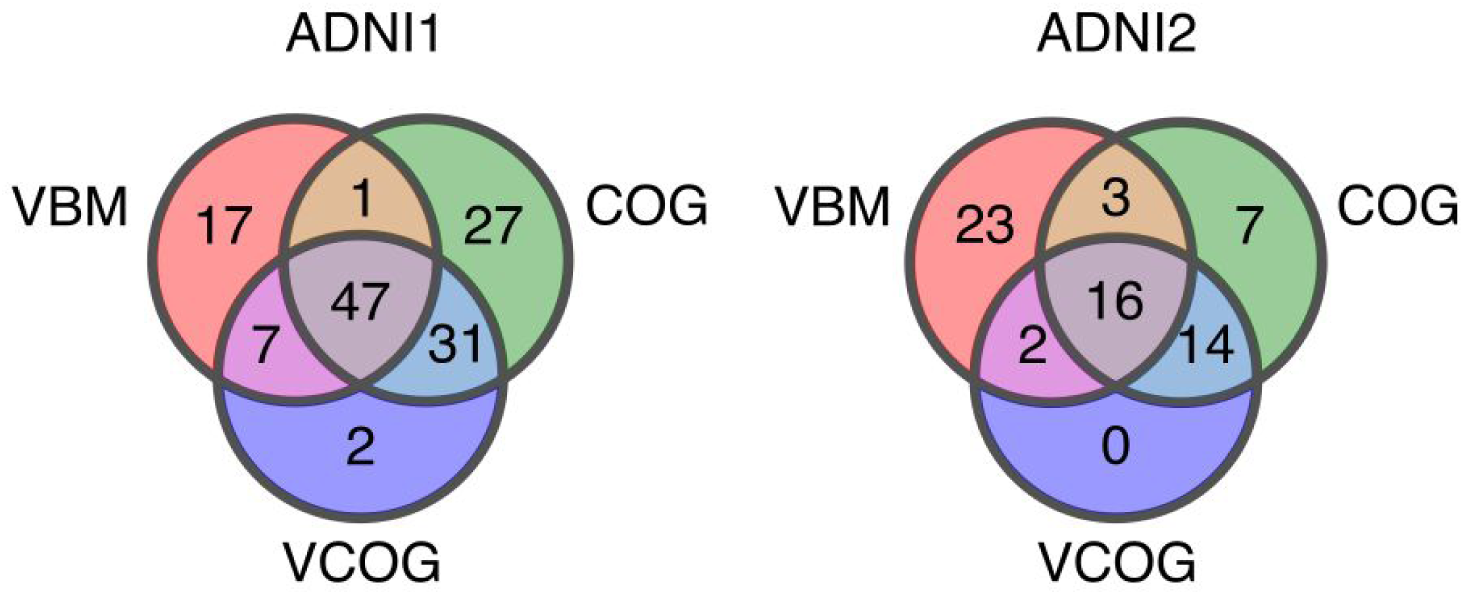
Venn diagram depicting the number of MCI subjects labeled as high confidence by the VBM, COG, and VCOG HPS models in ADNI1 and ADNI2.

Out of the high confidence subjects labeled by all three signatures, 97.9% and 93.7% from ADNI1 and ADNI2 respectively progressed to dementia (Supplementary Table S2). These subjects had worse cognition based on the MMSE and higher proportions of APOE4 carriers, A*β* positive and tau positive individuals, compared to the baseline rates in all MCI subjects. Of the high confidence subjects who were labeled only by the VBM model, 70.6% and 43.4% from ADNI1 and ADNI2 respectively were progressors. This group of subjects had less A*β* and tau positive individuals compared to the baseline rates. Of the high confidence subjects who were labeled only the COG model, 70.4% and 57.1% from ADNI1 and ADNI2 respectively progressed to dementia. This group appeared to have a greater proportion of A*β* positive individuals compared to the baseline rates in both ADNI1 and ADNI2 cohorts. The majority of these COG high confidence subjects were also male. Given the distinct characteristics among the exclusively COG, exclusively VBM, and VCOG high confidence subjects, these groups may represent subgroups with different risks for AD dementia. As it appears that a greater proportion of pMCI is captured when cognitive and structural MRI features are combined, these findings may support joining multiple modalities together in order to achieve higher positive predictive value. However, these results are qualitative and of an exploratory nature due to low sample sizes.

## Discussion

We developed a MRI and cognitive-based model to predict AD dementia with high PPV and specificity. Specifically, our two-stage predictive model reached 93.2% specificity and 93.1% PPV (80.4% when adjusted for 33.6% prevalence of progressors) in ADNI1 when classifying progressor vs stable MCI patients (within 3 years follow-up). We replicated these results in ADNI2 where the model reached 96.7% specificity and 81.2% PPV (87.8% adjusted PPV). With respect to specificity and PPV, these results are a substantial improvement over previous works combining structural MRI and cognition on the same prediction task, that have reported up to 76% specificity and 65% PPV (adjusted for 33.6% prevalence of progressors) [7]. Finally, our results also reproduced our past work which developed a model that optimizes specificity and PPV [20]. However, it appears that a combination of structural and functional MRI measures may lead to an improved prediction as two studies have reported 90-100% PPV with these measures [9,20], with the limitation of smaller sample sizes (56 total MCI subjects in [20], 86 total MCI subjects in [9]) due to the limited availability of functional MRI data in ADNI. Our proposed signature is based on widely available measures, and can be readily tested in many clinical trials. Functional MRI measures, by contrast, are only gaining traction in large clinical studies, and will at the minimum require more time to get widely adopted, if the very high PPVs are replicated in larger samples.

An ideal model to predict conversion to AD dementia would have both high sensitivity and specificity. However, the pathophysiological heterogeneity of clinical diagnosis will prevent highly accurate prediction linking brain features to clinical trajectories. We argue that, faced with heterogeneity, it is necessary to sacrifice sensitivity to focus on a subgroup of individuals with similar brain abnormalities. Due to the expected trade-off between specificity and sensitivity, the high specificity of our two-stage model indeed came at a cost of reduced sensitivity (55.1% in ADNI1 and 47.3% in ADNI2 for classifying pMCI vs sMCI), which is much lower than sensitivity values of 64%-95% reported by other groups [7,8,10–16]. The two-stage procedure did not offer gains compared to a simpler SVM model, if the threshold of the SVM model could be selected *a priori* to match the specificity of the two-stage procedure (see ROC curves in Figure 3). The two-stage prediction model offered the advantage of a principled approach to train the prediction model in order to maximize specificity, based on samples that are robust and easily classifiable, without testing a range of prediction thresholds. The choice of a L1 regularized logistic regression also led to a compact and interpretable subset of features for the HPS.

Favoring specificity over sensitivity is useful in settings where false positives need to be minimized and PPV needs to be high, such as expensive clinical trials. Here, with our VCOG HPS model, we report the highest PPVs for progression to AD from the MCI stage (up to 87.8%, adjusted for 33.6% prevalence of progressors) for models that included structural MRI and cognitive features, which are, importantly, modalities that are already widely used by clinicians. The present work could be used as a screening tool for recruitment in clinical trials that target MCI subjects who are likely to progress to dementia within three years. The implementation of an automated selection algorithm could also result in groups of MCI subjects with more homogeneous brain pathology. However, we note that high confidence subjects did not all present with significant amyloid burden (92.0% and 68.4% of high confidence subjects in ADNI1 and ADNI2 respectively, Figure 4), which means that not all high confidence individuals are likely to have prodromal AD, even when progressing to dementia.

When we trained our model with cognitive features only, tests for general cognition, memory, and executive function were chosen as the strongest predictors of AD dementia. Our COG HPS model thus supports previous research that reported general cognition, memory, and executive function as important neuropsychological predictors of dementia [7,29,38,39]. Compared to the state-of-the-art multi-domain cognition-based predictive model, which reported 87.1% specificity and 81.8% PPV (77.5% when adjusted to 33.6% pMCI prevalence) [40], our COG HPS model achieved similar performance reaching between 87.5%-95% specificity and 72.3%-85.1% (adjusted) PPV. As general cognition was the strongest feature in our model to predict progression, this supports previous findings that MCI patients with deficits across multiple domains are at the highest risk for dementia [39,41].

For our VBM model, we extracted a number of gray matter atrophy subtypes that recapitulated previously reported subtypes, namely the medial temporal lobe and parietal dominant subtypes [5,17–19], which were associated strongly with a diagnosis of AD dementia. Weights for the parietal dominant and medial temporal lobe subtypes (Subtypes 1 and 6 from Figure 1b, respectively) contributed substantially to the highly predictive signature in the VBM model. The atrophy pattern of subtype 6 is spatially similar to the spread of neurofibrillary tangles in Braak stages III and IV [42], which may support previous findings that tau aggregation mediates neurodegeneration [43]. The contributions of the parietal dominant and medial temporal lobe subtypes in the VBM HPS model are also in line with previous works, which have reported that cortical thickness and volumes of the medial temporal lobes, inferior parietal cortex, and precuneus are strong predictors of progression to dementia [7,11].

When combined with cognitive tests in the VCOG model, the structural subtypes were given marginal weights. This suggests some redundancy between atrophy and cognition, and that cognitive features have higher predictive power than structural features in the ADNI MCI sample. This conclusion is consistent with the observation that the COG model significantly outperformed the VBM model, similar to previous work [7]. Although cognitive markers were stronger features, the VCOG model assigned large negative weights for the structural subtypes 3, which showed greater relative GMV in the temporal lobes, and 7, which showed greater relative GMV in the parietal, occipital, and temporal lobes. This means that these features were predictive of stable MCI in the VCOG model, in line with previous work showing that atrophy in these regions is predictive of progression to dementia [7,11]. Furthermore, we demonstrated that combining MRI data with cognitive markers significantly improves upon a model based on MRI features alone. This result is again in line with the literature [7,10], yet was shown for the first time for a model specifically trained for high PPV. Note that in the current study, the predictive model was trained exclusively on images acquired on 1.5T scanners from ADNI1. Good generalization to ADNI2 with 3T scanners demonstrates robustness of imaging structural subtypes across scanner makes.

The VCOG highly predictive signature might reflect a late disease stage. We looked at the ratio of early to late MCI subjects in the ADNI2 sample (note that ADNI1 did not have early MCI subjects). Of the MCI subjects identified as high confidence by the VCOG model, 84.4% were late MCI subjects, compared to a rate of 34.9% of late MCI subjects in the entire ADNI2 MCI sample (Supplementary Figure S1). This approach may not be optimal for early detection of future cognitive decline. Training a model to classify MCI progressors and non-progressors to dementia could be done in order to capture future progressors in earlier preclinical stages (e.g. early MCI). Finally, we focused on structural MRI and neuropsychological batteries as features in our models due to their wide availability and established status as clinical tools. However, we believe adding other modalities such as PET imaging, CSF markers, functional MRI, genetic factors, or lifestyle factors could result in higher predictive power, especially at earlier preclinical stages of AD.

## Conclusion

In summary, we found a subgroup of patients with MCI who share a signature of cognitive deficits and brain atrophy, that put them at very high risk to progress from MCI to AD dementia within a time span of three years. We validated the signature in two separate cohorts that contained both stable MCI patients and MCI patients who progressed to dementia. The model was able to predict progression to dementia in MCI patients with up to 93.1% PPV and up to 96.7% specificity. The signature was present in about half of all progressors, demonstrating that gains in PPV can be made by focusing on a homogeneous, yet relatively common subgroup. Our model could potentially improve subject selection in clinical trials and identify individuals at a higher risk of AD dementia for early intervention in clinical settings.

## Acknowledgments

We thank Sylvia Villeneuve, Alexa Pichet-Binette and Jacob Vogel for providing us with data to help start our preliminary analyses for this project. We thank Hien Nguyen for advising us on parts of the statistical analyses.

Data collection and sharing for this project was funded by the Alzheimer’s Disease Neuroimaging Initiative (ADNI) (National Institutes of Health Grant U01 AG024904) and DOD ADNI (Department of Defense award number W81XWH-12-2-0012). ADNI is funded by the National Institute on Aging, the National Institute of Biomedical Imaging and Bioengineering, and through generous contributions from the following: AbbVie, Alzheimer’s Association; Alzheimer’s Drug Discovery Foundation; Araclon Biotech; BioClinica, Inc.; Biogen; Bristol-Myers Squibb Company; CereSpir, Inc.; Cogstate; Eisai Inc.; Elan Pharmaceuticals, Inc.; Eli Lilly and Company; EuroImmun; F. Hoffmann-La Roche Ltd and its affiliated company Genentech, Inc.; Fujirebio; GE Healthcare; IXICO Ltd.; Janssen Alzheimer Immunotherapy Research & Development, LLC.; Johnson & Johnson Pharmaceutical Research & Development LLC.; Lumosity; Lundbeck; Merck & Co., Inc.; Meso Scale Diagnostics, LLC.; NeuroRx Research; Neurotrack Technologies; Novartis Pharmaceuticals Corporation; Pfizer Inc.; Piramal Imaging; Servier; Takeda Pharmaceutical Company; and Transition Therapeutics. The Canadian Institutes of Health Research is providing funds to support ADNI clinical sites in Canada. Private sector contributions are facilitated by the Foundation for the National Institutes of Health (www.fnih.org). The grantee organization is the Northern California Institute for Research and Education, and the study is coordinated by the Alzheimer’s Therapeutic Research Institute at the University of Southern California. ADNI data are disseminated by the Laboratory for Neuro Imaging at the University of Southern California.

The computational resources used to perform the data analysis were provided by Compute Canada (www.computecanada.org). This project was funded by NSERC grant number RN000028 and the Canadian Consortium on Neurodegeneration in Aging (CCNA, www.ccna-ccnv.ca), through a grant from the Canadian Institutes of Health Research and funding from several partners including SANOFI-ADVENTIS R&D. AT was supported by a bursary from the Centre de recherche de l’institut universitaire de gériatrie de Montréal and the Courtois foundation. CD was supported by a salary award from the Lemaire foundation and Courtois foundation. PB was supported by a salary award from Fonds de recherche du Québec -- Santé and the Courtois foundation.

