## Supplemental material for "A highly predictive signature of cognition and brain atrophy for progression to Alzheimer’s dementia"

### Supplementary methods

**Multiple machine learning algorithms.** In addition to the support vector machine with a linear kernel and the two-stage highly predictive signature (HPS), we applied other machine learning algorithms to train models to distinguish between AD patients and CN subjects in the ADNI1 dataset. These latter algorithms included a support vector machine (SVM) with a radial basis function kernel, K nearest neighbors, random forest, and Gaussian naive Bayes. We used a nested cross-validation loop (stratified shuffle split with 50 splits and 20% test size) in our grid searches for optimal hyperparameters for the SVM, K nearest neighbors, and random forest algorithms. After training on AD vs CN in ADNI1, we validated each model on classifying pMCI vs sMCI in ADNI1. We then tested the model performance on ADNI2 to classify AD vs CN and then pMCI vs sMCI. All algorithms were implemented by Scikit-learn (1). See below for details about hyperparameter optimization for the SVM, K nearest neighbors, and random forest.

**SVM with radial basis function kernel.** We optimized for the hyperparameter C (grid was $10^{-2}$ to $10^{1}$ with 15 equal steps).

**K nearest neighbors.** The grid search for optimal hyperparameters included four different values of K (3, 4, 5, 6) and two weight functions used for the prediction ("uniform" where all points in the neighborhood are weighted equally, and "distance" where points are weighted inversely to their distance).

**Random forest.** Our grid search included number of trees (10 or 25), number of maximum features to consider for the best split (5 or all possible features (VBM: 11, COG: 9, VCOG: 16)), maximum depth of tree (10, 50 or unlimited), and the option to bootstrap the samples when building the trees.

### Supplementary figures

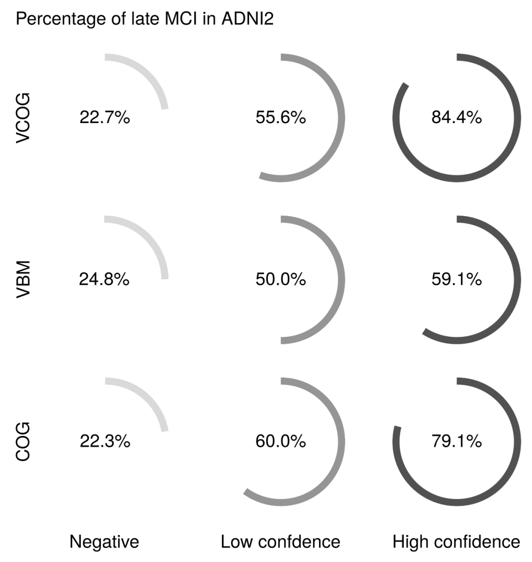


**Fig. S1.** Percentage of late MCI ADNI2 subjects within each HPS grouping (high confidence, low confidence, negative) across each highly predictive model (VCOG, VBM, COG). In each model, there was a significantly greater proportion of late MCI subjects in the high confidence class compared to the negative class (VCOG: $\chi^2 = 51.0, p < 0.001$; VBM: $\chi^2 = 21.8, p < 0.001$; COG: $\chi^2 = 59.9, p < 0.001$).

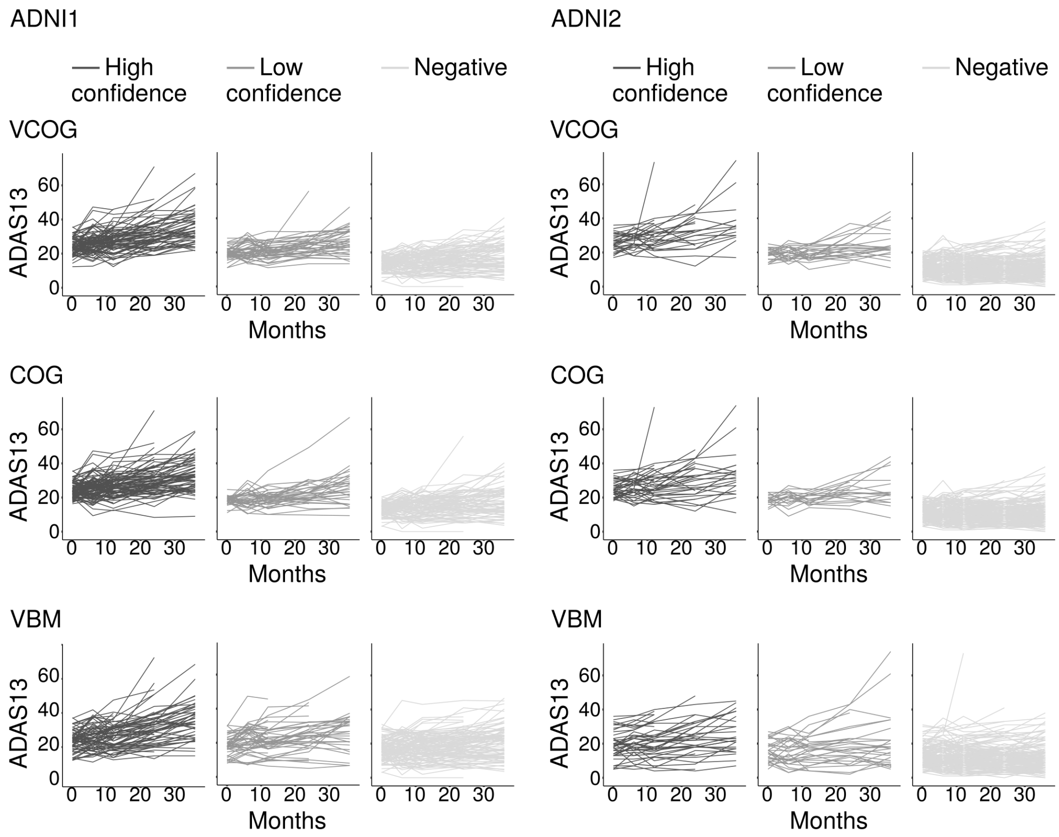


**Fig. S2.** Cognitive trajectories for individual MCI subjects in ADNI1 and ADNI2 grouped by HPS classifications (high confidence, low confidence, negative) by the VCOG, COG, and VBM high confidence prediction models.

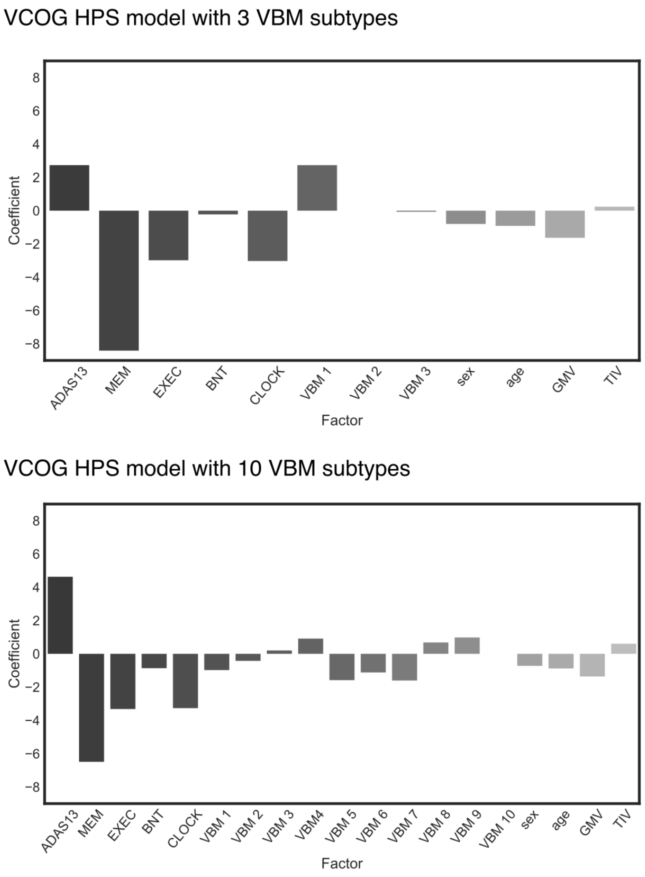


**Fig. S3.** Coefficients of factors in the VCOG HPS models for a model featuring 3 VBM subtypes and a model featuring 10 VBM subtypes.

### Supplementary tables

**Table S1. pMCI vs sMCI performance metrics for VCOG HPS models with different number of VBM subtypes.**

| **ADNI1** | **3 clusters** | **7 clusters** | **10 clusters** |
|---|---|---|---|
| Specificity | 0.8636 | 0.9310 | 0.8864 |
| Sensitivity | 0.5578 | 0.5510 | 0.5510 |
| Positive predictive value (adjusted) | 0.6743 | 0.8035 | 0.7105 |
| Accuracy | 0.6723 | 0.6936 | 0.6766 |
| **ADNI2** | **3 clusters** | **7 clusters** | **10 clusters** |
| Specificity | 0.9556 | 0.9667 | 0.9556 |
| Sensitivity | 0.4727 | 0.4727 | 0.4364 |
| Positive predictive value (adjusted) | 0.8433 | 0.8777 | 0.8324 |
| Accuracy | 0.8425 | 0.8511 | 0.8340 |

Positive predictive value was adjusted for a 33.6% prevalence of pMCI subjects.

**Table S2. Characteristics of high confidence subjects from the VBM, COG, and VCOG signatures in ADNI1 and ADNI2 MCI cohorts.**

| **ADNI1** | **Baseline** | **VBM**<br>~~COG~~<br>~~VCOG~~ | ~~VBM~~<br>**COG**<br>~~VCOG~~ | ~~VBM~~<br>~~COG~~<br>**VCOG** | **VBM**<br>**COG**<br>~~VCOG~~ | **VBM**<br>~~COG~~<br>**VCOG** | ~~VBM~~<br>**COG**<br>**VCOG** | **VBM**<br>**COG**<br>**VCOG** |
|---|---|---|---|---|---|---|---|---|
| pMCI % | 62.6 | 70.6 | 70.4 | 100 | 100 | 57.1 | 93.5 | 97.9 |
| APOE4 % | 57.0 | 58.8 | 70.4 | 100 | 100 | 71.4 | 71.0 | 63.8 |
| A$\beta$+ % | 76.1 | 50.0 | 91.7 | n/a | 100 | 100 | 88.9 | 91.7 |
| tau+ % | 50.7 | 0.0 | 83.3 | n/a | 0.0 | 50.0 | 66.7 | 66.7 |
| Female % | 40.8 | 41.2 | 29.6 | 50.0 | 0.0 | 42.9 | 61.3 | 36.2 |
| Age | 74.1±7.2 | 74.4±5.5 | 73.9±6.0 | 80.0±6.2 | 79.3 | 69.3±4.1 | 73.9±7.0 | 74.4±7.3 |
| Education | 15.7±2.9 | 17.1±2.1 | 15.7±3.3 | 15.0±1.4 | 20.0 | 16.0±2.6 | 14.3±3.1 | 15.9±2.9 |
| MMSE | 27.0±1.8 | 27.8±1.4 | 27.6±1.2 | 26.5±0.7 | 29.0 | 25.4±0.5 | 25.9±1.5 | 26.0±1.6 |
| **ADNI2** | **Baseline** | **VBM**<br>~~COG~~<br>~~VCOG~~ | ~~VBM~~<br>**COG**<br>~~VCOG~~ | ~~VBM~~<br>~~COG~~<br>**VCOG** | **VBM**<br>**COG**<br>~~VCOG~~ | **VBM**<br>~~COG~~<br>**VCOG** | ~~VBM~~<br>**COG**<br>**VCOG** | **VBM**<br>**COG**<br>**VCOG** |
| pMCI % | 30.6 | 43.5 | 57.1 | n/a | 33.3 | 0.0 | 78.6 | 93.7 |
| APOE4 % | 42.6 | 47.8 | 42.9 | n/a | 33.3 | 50.0 | 57.1 | 68.7 |
| A$\beta$+ % | 69.0 | 75.0 | 100 | n/a | 100 | 100 | 55.6 | 77.8 |
| tau+ % | 39.7 | 42.9 | 25.0 | n/a | 50.0 | 0.0 | 62.5 | 55.6 |
| Female % | 48.1 | 39.1 | 14.3 | n/a | 33.3 | 50.0 | 57.1 | 50.0 |
| Age | 70.7±7.3 | 71.4±5.9 | 74.8±6.6 | n/a | 67.0±13.7 | 73.6±2.5 | 71.7±8.6 | 77.1±4.8 |
| Education | 16.4±2.6 | 16.3±2.6 | 16.9±2.5 | n/a | 15.0±3.6 | 19.0±1.4 | 15.6±2.3 | 15.9±2.8 |
| MMSE | 28.2±1.7 | 28.3±1.5 | 28.0±1.3 | n/a | 28.7±1.5 | 27.5±2.1 | 26.7±1.8 | 26.1±1.7 |

The Baseline column represents values for all MCI subjects.
Age and education are presented in years (mean ± standard deviation).
A$\beta$ and tau CSF measures were available for approximately one third of all subjects across both cohorts.
